# Transcriptomic analysis of cecal mucosal immunity in SPF White Leghorn chicks infected with precocious and parent strains of *Eimeria tenella*

**DOI:** 10.64898/2026.07.24.740658

**Authors:** Wenyu Ma, Kexin Du, Tingting Yi, Xuefei Liang, Shuanjing Niu, Xuewei Liu, Mengze Du, Jian An, Deqi Yin, Qingming Li

## Abstract

*Eimeria tenella* (*E. tenella*) preferentially invades the cecum of young chickens and causes enormous economic losses to the global poultry industry. In this study, chick infection models of virulent parent strain and precocious attenuated line were established with schizogony (2 dpi) and gametogony (6 dpi) as two critical sampling time points. Combined with pathogenicity detection, transcriptome sequencing, mucosal immune index measurement and homologous challenge protection assays, we systematically deciphered the differential molecular mechanisms underlying pathogenicity and immune regulation between the two strains. Pathogenicity results showed that increased infection dosages suppressed weight gain and aggravated bloody diarrhea and oocyst shedding in both strains. In particular, infection with 1 x 10^5^ sporulated oocysts of the parent strain caused massive chick mortality, while the precocious line exhibited markedly lower virulence. Transcriptomic data revealed that gametogony (6 dpi) represented the peak of host immune response. The parent strain persistently overactivated the NF- κB - mediated positive feedback cascade of coagulation and complement as well as ECM remodeling, triggering steroid metabolic disorder and antioxidant exhaustion, which ultimately induced severe hemorrhagic necrosis of the cecum. By contrast, the precocious line specifically activated the PPAR - γ signaling pathway to negatively restrain excessive inflammation, accompanied by enriched TLR signaling and leukocyte transendothelial migration pathways, thereby forming an immune cascade of "lipid anti - inflammation - pathogen elimination - mucosal repair". Immunoprotection trials verified that priming with 1 × 10^4^ sporulated oocysts of the precocious line significantly alleviated cecal lesions and reduced oocyst output upon secondary challenge, conferring stable mucosal immune protection. The expression trends of seven hub genes (*PPARG, PLIN1, CYP1A1, THBS1, FMO4, CYP2C18, CYP14*) detected via qRT - PCR were consistent with RNA - seq data. This study identified a dual regulatory paradigm consisting of NF - κB - mediated tissue injury and PPAR - γ - dependent anti - inflammatory responses, refined the mucosal immune theoretical framework for commercially available precocious attenuated strains, and provided candidate molecular targets for targeted anti - coccidial intervention in chickens.

## Introduction

Avian coccidiosis is an intestinal parasitic disease caused by protozoans of the genus *Eimeria* (Nguyen et al., 2026). Among all chicken coccidia, *Eimeria tenella* (*E. tenella*) exerts the highest pathogenicity in chicks aged 15 to 50 days (Liu et al., 2024). Coccidiosis incurs annual economic losses exceeding 14 billion US dollars worldwide, posing a severe threat to the sustainable development of the poultry industry (Bafundo, 2025). The parasite transmits mainly via the fecal - oral route, with lesions specifically localized in the cecum. Clinically infected chicks manifest reduced feed intake, intestinal inflammation, hemorrhagic diarrhea and growth retardation; severe infection may result in extensive cecal necrosis and even massive chicken death (Ewais et al., 2024). Although live attenuated vaccines derived from precocious lines have been commercially produced, the molecular regulatory network of host mucosal immunity responsible for their low virulence and high immunogenicity remains poorly characterized.

High - throughput RNA - seq provides a powerful tool to dissect the molecular crosstalk between coccidian parasites and their avian hosts, and multi-omics research focusing on *E. tenella* has expanded rapidly in recent years (Ferreira et al., 2023). Previous transcriptomic analyses indicated that overexpression of malate dehydrogenase is closely associated with parasite endogenous development, host cell invasion and drug resistance formation (Yue et al., 2010). Single - cell RNA sequencing demonstrated that *E. tenella* infection drives prominent epithelial apoptosis, accompanied by reduced abundance of cell subsets maintaining mitochondrial and cytoplasmic homeostasis (Tu et al., 2025). Widespread dynamic m^7^G mRNA methylation during oocyst sporulation was further uncovered. This epigenetic modification modulates the transcription of metabolic and developmental genes and directly determines oocyst infectivity (Fan et al., 2024). Existing comparative transcriptomics between precocious lines and parent strains solely focused on merozoite - derived virulence genes such as microneme, rhoptry and dense granule proteins (Zhang, 2020). These parasite - centric investigations fail to explain the distinct phenotypic differences: mild intestinal damage and superior protective immunity induced by precocious line infection. Current relevant research bears prominent limitations. Most omics studies only adopt a single infection time point, lacking temporal comparisons of host cecal mucosal immunity across schizogony and gametogony. Moreover, transcriptomic profiling of precocious lines largely concentrates on parasite genes while neglecting dynamic host immune remodeling along parasite developmental stages. Schizogony at 2 dpi and gametogony at 6 dpi constitute two critical time windows determining strain virulence and immunogenicity. To date, no research has simultaneously compared host transcriptional profiles induced by parent and precocious strains at these two key stages, creating a critical knowledge gap hindering in - depth interpretation of precise anti - coccidial strategies. In the present study, we systematically evaluated the in vivo pathogenic differences between parent and precocious *E. tenella* strains, employed RNA - seq to compare cecal transcriptional profiles at 2 dpi and 6 dpi post-infection, and elucidated the host molecular mechanisms driving divergent pathogenicity, offering theoretical evidence for deciphering the immunological basis of commercial precocious live vaccines and screening novel targeted anti - coccidial molecular markers.

## Materials and methods

### Ethical statement

All animal procedures in this experiment were reviewed and approved by the Institutional Animal Welfare and Ethics Committee of Beijing University of Agriculture (Ethics approval No. BUA2022080). All animal procedures were performed in accordance with the regulations and guidelines of the committee to ensure humane treatment of chicks.

### Animals

A total of 150 one - day - old male SPF White Leghorn chicks were purchased from Beijing Boehringer Ingelheim Vital Animal Biotechnology Co., Ltd. The chicks were raised at a constant temperature of 22 ± 2 ℃ under a 12 h light/12 h dark photoperiod, and fed with commercial coccidia-free and antibiotic-free complete chick feed (XTC07YJ-001, Jiangsu, China) throughout the feeding period.

### Eimeria tenella oocysts

The virulent parent Henan strain and its derived precocious line of *E. tenella* were preserved in the Chicken Coccidiosis Laboratory, College of Animal Science and Technology, Beijing University of Agriculture. Oocysts were propagated one month prior to experiments, sporulated in 2.5% potassium dichromate solution, and stored at 4℃ for subsequent use. The prepatent period of the parent strain was 143 h, while repeated chicken passages shortened the prepatent period of the precocious line to 121 h.

### Experimental design

As illustrated in Fig. 1A, sixty - three healthy 28 - day - old chicks with body weight variations within ± 5 g were randomly divided into seven groups with nine birds per group, including a negative control group (NC), parent strain infection groups (E.ten), and precocious line infection groups (Pre*E. ten*). At 28 days of age, chicks in infection groups were orally inoculated with 1 × 10^3^, 1 × 10^4^ or 1 × 10^5^ sporulated oocysts of the parent strain or precocious line for pathogenicity assessment. At 2 and 6 dpi, three chicks from the NC group and each 1 × 10^4^ infection subgroup were sacrificed to collect cecal tissues for histopathological section preparation. At 7 dpi, body weight gain, mortality, bloody stool scores and oocysts per gram of cecal contents (OPG) were recorded.

**Fig. 1.**
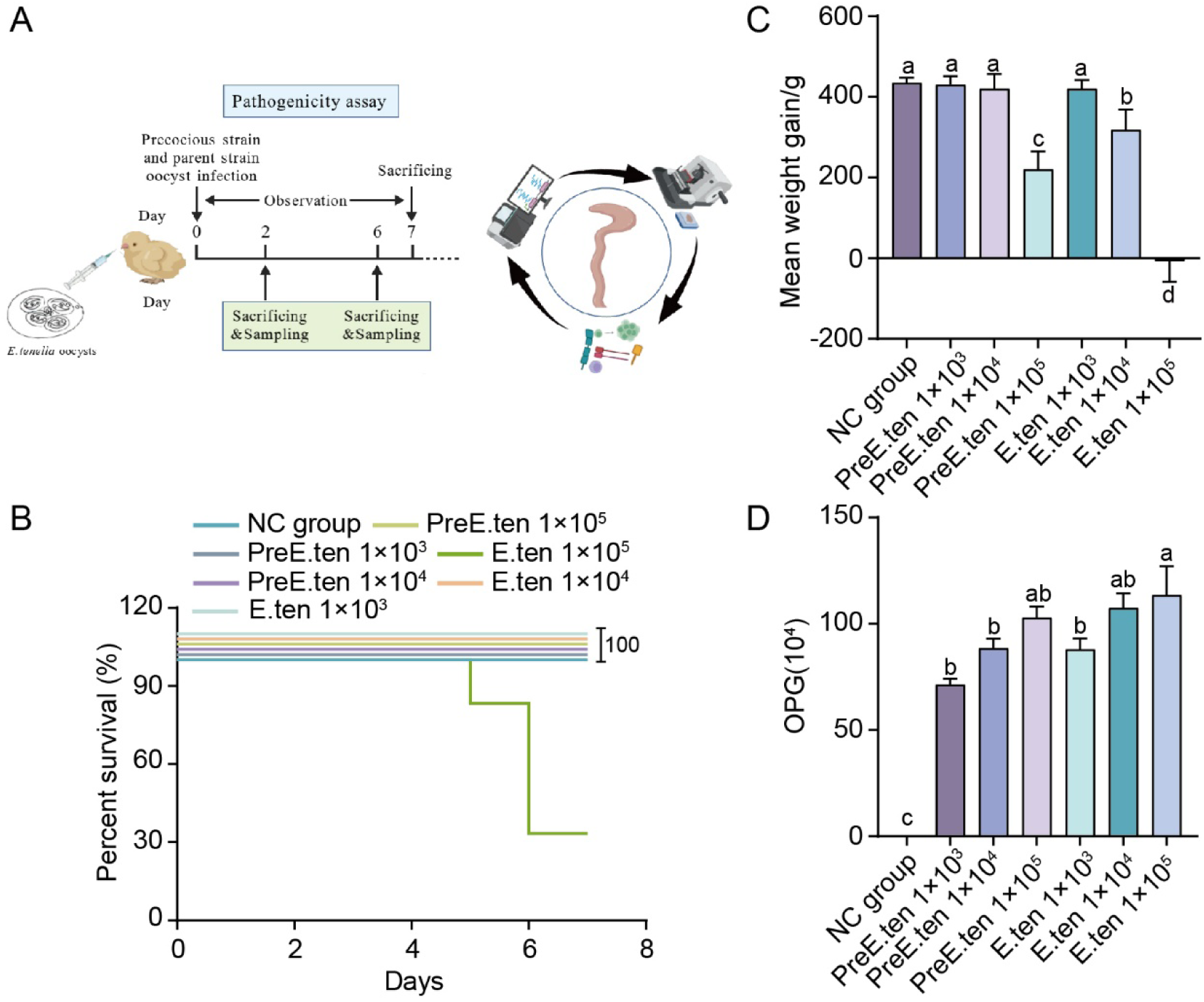
Pathogenicity detection of sporulated oocysts of *E. tenella* precocious line and parent strain. (A) Schematic diagram of the experimental design. (B) Effects of different dosages of precocious line and parent strain oocysts on the survival rate of chicks during pathogenicity detection. (C) Effects of different dosages of precocious line and parent strain oocysts on body weight gain of chicks during pathogenicity detection. (D) OPG of cecal contents in each group. Values marked with different lowercase superscript letters (a-d) within the same column indicate significant differences at *P* < 0.05.

### Bloody stool scoring

Bloody stool conditions of each experimental group were observed daily from day 4 to day 7 post-infection. A scoring scale ranging from 0 to 4 was adopted in accordance with the method described by Youn. et al. (2001).

### Quantification of oocysts per gram of cecal contents

At 7 dpi, 2 g cecal content samples were collected from each group. Oocyst counting was performed using the McMaster technique as described by Joyner.et al. (1970).

### Measurement of immune organ indices

Chicks were weighed before euthanasia. The intact spleen, bursa of Fabricius and thymus were aseptically dissected, with excess fat and connective tissues removed using forceps and scissors. Fresh organ weights were recorded to calculate immune organ indices, which were defined as organ weight/live body weight.

### RNA extraction and transcriptome sequencing

Twenty - eight - day - old SPF chicks were randomly assigned to infection subgroups (n = 12) and the negative control subgroup (n = 3). Chicks in infection groups received an oral dose of 1 × 10^4^ sporulated oocysts of either the precocious line or parent strain, whereas control chicks were administered an equal volume of sterile normal saline. Segments of cecal tissue (2 cm in length) were harvested at 2 and 6 dpi, immersed in Trizol reagent and preserved in liquid nitrogen. Library construction and paired - end sequencing were conducted on the HiSeq platform by Shanghai Bioengineering Co., Ltd. Sequencing quality control thresholds were set as Q20 ≥ 95% and Q30 ≥ 85%. Differentially expressed genes (DEGs) were screened with the criteria of ∣log2FC∣ > 2 and *P* < 0.05, followed by functional enrichment analyses of Gene Ontology (GO) and Kyoto Encyclopedia of Genes and Genomes (KEGG) to identify significantly enriched GO terms and signaling pathways. Statistical significance was defined as *P* < 0.05. All raw sequencing reads generated in this study have been deposited in the NCBI Sequence Read Archive (SRA) under accession number PRJNA1491229 (https://www.ncbi.nlm.nih.gov/sra/PRJNA1491229)

### Validation of RNA-seq data via quantitative real-time PCR

Quantitative real-time PCR was performed to verify the mRNA expression levels of seven candidate genes (*PPARG, PLIN1, CYP1A1, THBS1, FMO4, CYP2C18, CXCL14*) identified from transcriptome data, with β-actin used as the reference gene. The amplification program was set as follows: initial denaturation at 95℃ for 30 s, followed by 40 cycles of 95℃ for 15 s and 60℃ for 30 s, with a melting curve generated from 65℃ to 95℃. The relative gene expression was calculated using the 2^-ΔΔCt^ method. Primer sequences are listed in Table 1. All qRT-PCR reactions were conducted in three technical replicates.

**Table 1.**
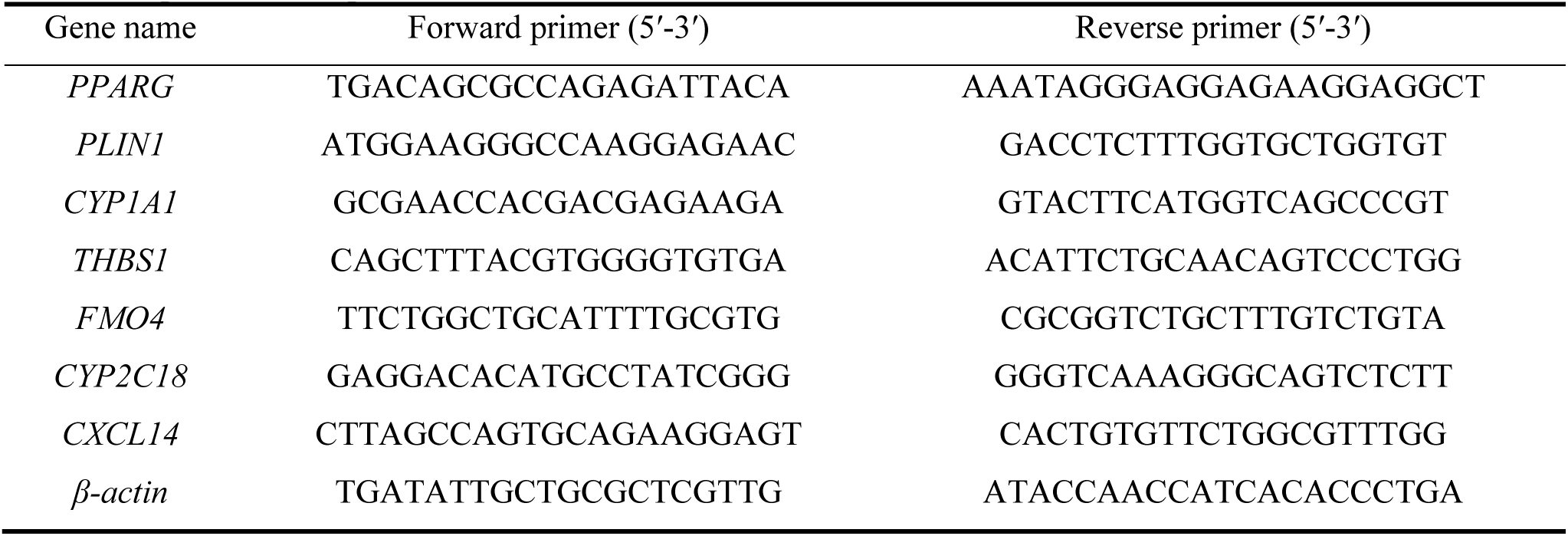
Primer sequences for qRT-PCR.

### Histopathological analysis

Cecal tissues (5 cm long) were collected at 2 and 6 dpi, fixed in 4% paraformaldehyde (Biosharp, Beijing, China) for 72 h, processed into paraffin sections and stained with hematoxylin and eosin for microscopic observation. Remaining cecal tissues were preserved for subsequent immune functional assays.

### Isolation and proliferation assay of cecal lymphocytes

Cecal tissues were harvested under aseptic conditions, minced, homogenized, and subjected to density gradient centrifugation to isolate lymphocytes. After washing, cells were resuspended for subsequent experiments. A total of 2 × 10^5^ lymphocytes suspended in 200 μL medium were seeded into each well of a 96 - well plate, followed by stimulation with ConA (Biosharp, Beijing, China) for 72 h. Optical density at 490 nm was detected according to the manufacturer’s instructions of the MTS kit(Promega, Beijing, China), and the stimulation index (SI) was calculated accordingly.

### Enzyme-linked immunosorbent assay

Cecal tissues were homogenized in nine volumes of pre - cooled PBS, and homogenates were centrifuged at 5000 rpm for 15 min at 4 ℃ to collect supernatants. The concentrations of secretory immunoglobulin A (SIgA) in cecal contents were determined using a chicken SIgA ELISA kit (mlbio, Shanghai, China), with absorbance measured at 450 nm.

### Immunoprotection assay

Thirty - six healthy 28 - day - old chicks with body weight variations within ± 5 g were randomly allocated into six groups with six individuals per group: negative control group (NC), parent strain priming groups (*E. ten*), precocious line priming groups (*PreE. ten*), and positive challenge control group (PC). At 28 days of age, primed chicks were orally inoculated with 1 × 10^3^ or 1 × 10^4^ sporulated oocysts of the parent strain or precocious line. At 42 days of age, all primed groups received a homologous challenge of 1 × 10^5^ sporulated oocysts of the parent strain. The PC group received no prior priming and was only challenged with 1 × 10^5^ parent strain oocysts at 42 days of age (Table 2). Detection indicators were consistent with those described in Section 2.4, and cecal OPG as well as immune organ indices were measured at 49 dpi.

**Table 2.**
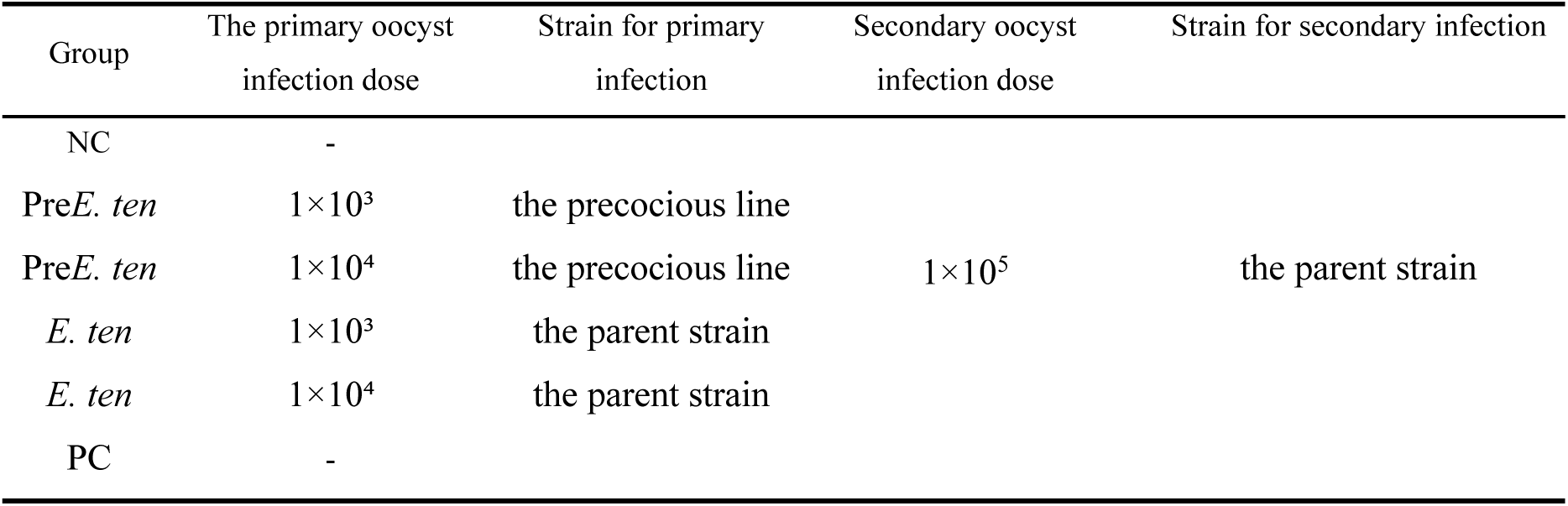
Infection protocol for immune protection assay.

### Statistical analysis

All experiments were performed in three independent biological replicates. Statistical analyses were carried out using GraphPad Prism 10.0 and SPSS 17.0 software. All data were presented as mean ± standard error (SE). Two - tailed unpaired Student’s t - test was used for comparisons between two groups, while one-way analysis of variance (ANOVA) was applied for multiple group comparisons. Values within the same column marked with different superscript lowercase letters (a - d) indicate significant differences at *P* < 0.05. Statistical significance levels were set as \*\**P* < 0.01 and \*\*\**P* < 0.001.

## Results

### Pathogenicity detection

Significant pathogenic differences were observed between the parent strain and precocious line of *E. tenella*. As shown in Fig. 1B, mortality occurred in chicks infected with 1 × 10^5^ sporulated oocysts of the parent strain, with a final survival rate of 33.3%, whereas no deaths were recorded in all other experimental groups. The body weight changes of chicks are presented in Fig. 1C. Compared with the NC group, the 1 × 10^5^ Pre*E. ten* group, 1 × 10^4^ *E. ten* group and 1 × 10^5^ *E. ten* group exhibited significantly reduced body weight gain. Notably, infection with 1 × 10^5^ parent strain oocysts induced negative body weight growth in chicks. OPG quantification of fecal samples revealed that both parent and precocious line infection groups showed statistically higher OPG values than the NC group, and the OPG levels increased in a dose-dependent manner (Fig. 1D). Bloody stool scoring results demonstrated that chicks in the 1 × 10^4^ and 1 × 10^5^ dosage groups of both strains developed bloody stool starting at 4 dpi, and the total bloody stool scores increased with the elevated infection dosage (Table 3).

**Table 3.** Statistical analysis of bloody stool scores after primary coccidial infection in different groups.

| Group | Oocyst infection dose | 84 h | 93 h | 108 h | 117 h | 132 h | 141 h | 156 h | 165 h | Score of bloody feces |
| --- | --- | --- | --- | --- | --- | --- | --- | --- | --- | --- |
| NC | 0 | 0 | 0 | 0 | 0 | 0 | 0 | 0 | 0 | 0<sup>c</sup> |
| Pre*E. ten* | 1.0×10<sup>3</sup> | 0 | 0 | 0 | 0 | 0 | 0 | 0 | 0 | 0<sup>c</sup> |
| Pre*E. ten* | 1.0×10<sup>4</sup> | 2 | 3 | 1 | 0 | 0 | 0 | 0 | 0 | 0.75±1.1<sup>bc</sup> |
| Pre*E. ten* | 1.0×10<sup>5</sup> | 3 | 4 | 2 | 1 | 0 | 0 | 0 | 0 | 1.25±1.5<sup>ab</sup> |
| *E. ten* | 1.0×10<sup>3</sup> | 0 | 0 | 0 | 0 | 0 | 0 | 0 | 0 | 0<sup>c</sup> |
| *E. ten* | 1.0×10<sup>4</sup> | 0 | 1 | 3 | 2 | 1 | 0 | 0 | 0 | 0.87±1.1<sup>bc</sup> |
| *E. ten* | 1.0×10<sup>5</sup> | 0 | 2 | 4 | 4 | 3 | 2 | 1 | 1 | 2.13±1.4<sup>a</sup> |

Values marked with different lowercase superscript letters (a-d) within the same column indicate significant differences at $P < 0.05$ .

### Transcriptomic data analysis

Based on the pathogenicity results of chicks infected with different dosages and virulent *E. tenella* oocysts, the dosage of 1× 10^4^ sporulated oocysts was selected for subsequent transcriptomic analysis to further explore the molecular mucosal immune responses of chick ceca induced by precocious line and parent strain infections. Cecal tissues were collected from the blank control group (C group), precocious line-infected groups at 2 and 6 dpi (PreE2 and PreE6), and parent strain-infected groups at 2 and 6 dpi (E2 and E6) for transcriptome sequencing (Fig. 2A). A total of 15 cecal transcriptome libraries (3 biological replicates per group) were constructed and sequenced. The results yielded 36,376,946 to 58,106,532 paired-end raw reads (150 bp) per library, with GC contents ranging from 46.44% to 48.89%. Low-quality reads, ambiguous bases, and adapter sequences were filtered using Trimmomatic software, generating 36,332,048 to 58,033,254 valid clean reads with a total data volume of 97.9 Gb. The Q20 and Q30 values of all clean data ranged from 97.36% to 97.91% and 93.54% to 94.65%, respectively (Supplementary Tables S1 and S2). These quality control indicators confirmed that the transcriptomic data were reliable and qualified for subsequent analysis of cecal mucosal immune responses.

**Fig. 2.**
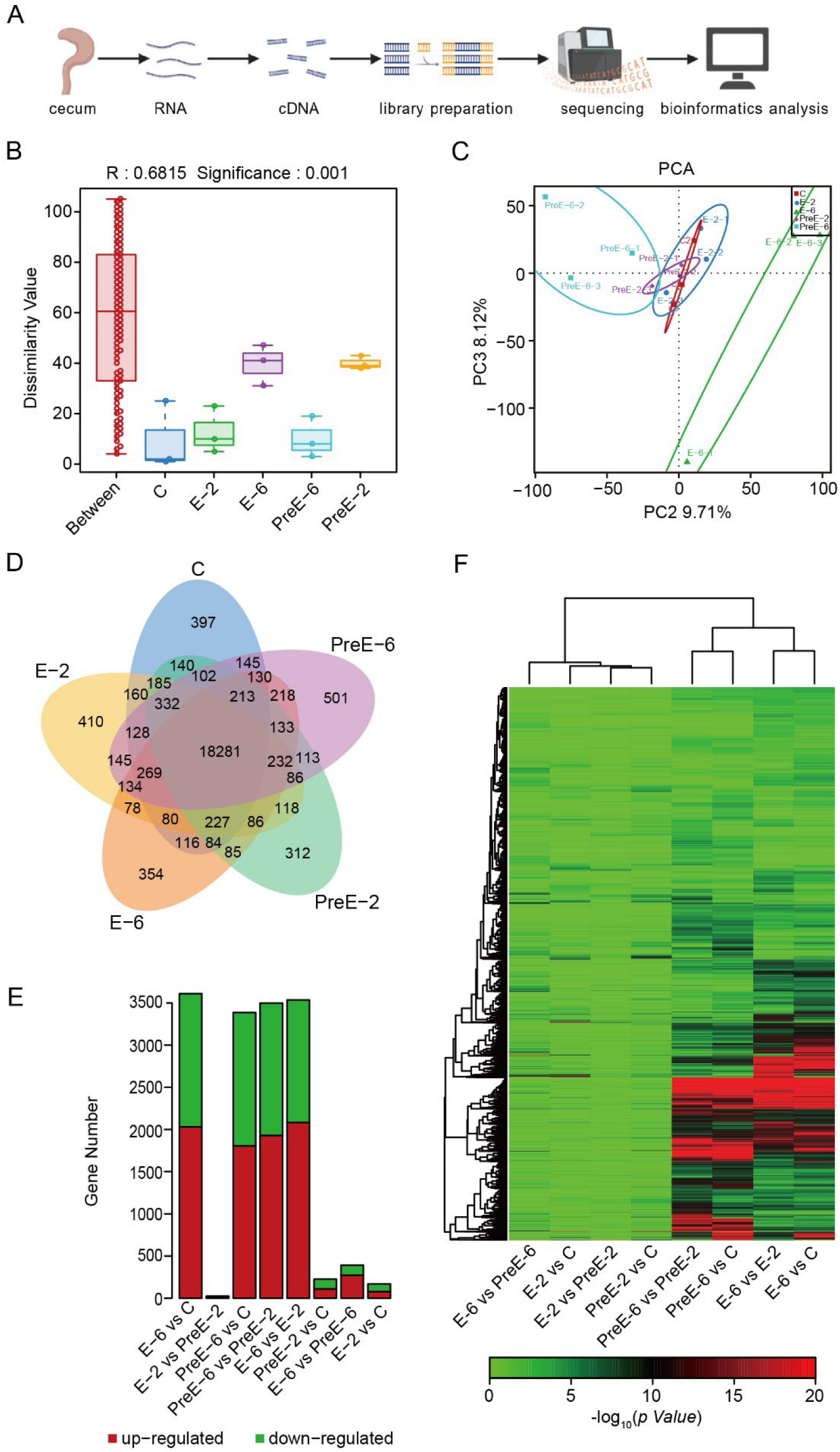
DEGs dentified among different groups. (A) Schematic workflow of transcriptome sequencing. (B) ANOSIM showing the discrepancies among groups. (C) PCA illustrating the overall differences across samples. (D) Venn diagram displaying the distribution of DEGs among groups. (E) Histogram presenting the numbers of significantly up and downregulated genes in different comparison groups. (F) Clustering heatmap of DEGs across different treatment groups.

ANOSIM showed that the R value was close to 1, indicating significantly higher intergroup differences than intragroup differences (Fig. 2B). PCA further revealed distinct transcriptional profiles among the C, PreE6, and E6 groups (Fig. 2C). Venn diagram analysis identified 18,281 conserved genes across all five groups. Specifically, the parent strain-infected groups (E2, E6) possessed 410 and 352 unique expressed genes, while the precocious line-infected groups (PreE2, PreE6) had 312 and 501 unique expressed genes, respectively, providing fundamental data for elucidating the differential cecal mucosal immune responses induced by the two virulent strains (Fig. 2D).

At the transcriptional level, host transcriptomic perturbation was mild at 2 dpi, whereas extensive immune transcriptional remodeling occurred at 6 dpi. Therefore, subsequent functional enrichment analyses were mainly focused on the 6 dpi samples. Differential expression analysis showed the following results: compared with the C group, the E6 group had 2,004 significantly upregulated genes and 1,555 significantly downregulated genes; the PreE6 group had 1,781 significantly upregulated genes and 1,557 significantly downregulated genes. Compared with the E2 group, the E6 group had 2,053 significantly upregulated genes and 1,432 significantly downregulated genes. Compared with the PreE2 group, the PreE6 group had 1,903 significantly upregulated genes and 1,547 significantly downregulated genes. In addition, the E2 group had 16 upregulated and 9 downregulated DEGs relative to the PreE2 group; the PreE2 group had 109 upregulated and 113 downregulated DEGs relative to the C group; the E2 group had 79 upregulated and 88 downregulated DEGs relative to the C group; and the E6 group had 271 upregulated and 116 downregulated DEGs relative to the PreE6 group (Fig. 2E). The clustering heatmap of common DEGs exhibited treatment- and time-specific clustering patterns of core DEGs across comparison groups, suggesting that 6 dpi serves as a critical regulatory node for host cecal mucosal immune responses triggered by *E. tenella* infection (Fig. 2F). Seven differentially expressed genes, including both upregulated and downregulated genes, were randomly selected from the transcriptomic dataset for quantitative real time PCR validation. As shown in Fig.S1, the expression trends of *PPARG*, *PLIN1*, *CYP1A1*, *THBS1*, *FMO4*, *CYP2C18*, and *CXCL14* were highly consistent with the RNA seq results, confirming the reliability and accuracy of the transcriptomic data obtained in this study.

### Transcriptomic analysis of host intestinal responses

Consistent with the DEG clustering heatmap results in Fig. 2F, both the precocious line and parent strain induced substantially stronger transcriptional responses at 6 dpi than at 2 dpi. Accordingly, subsequent functional analyses were focused on transcriptomic profiles at 6 dpi. GO enrichment analysis revealed that the DEGs identified from both precocious line comparisons (PreE6 vs C, PreE6 vs PreE2) and parent strain comparisons (E6 vs C, E6 vs E2) were predominantly enriched in cellular component terms including plasma membrane, membrane, and extracellular region. In terms of biological processes, DEGs from all comparison groups were commonly annotated to immune response, defense response, and cytokine production (Fig. 3A - D). Specifically, DEGs in the PreE6 group were prominently enriched in leukocyte activation and T cell activation, whereas DEGs in the E6 group were additionally enriched in cytokine mediated signaling pathways, cytokine responses, and cellular responses to cytokine stimulation.KEGG pathway enrichment further verified that 6 dpi represents a critical response node dominated by multiple core immune signaling pathways. Both PreE6 vs C and E6 vs C comparisons exhibited significant enrichment in Th17 cell differentiation, NF - κB signaling pathway, intestinal immune network for IgA production, complement and coagulation cascades, Th1/Th2 cell differentiation, and natural killer cell mediated cytotoxicity (Fig. 3E - F). Notably, the PreE6 group showed unique enrichment in the PPAR signaling pathway, chemokine signaling pathway, and leukocyte transendothelial migration pathway. In contrast, the E6 group was specifically enriched in steroid hormone biosynthesis, ECM receptor interaction, and fatty acid elongation pathways.

**Fig. 3.**
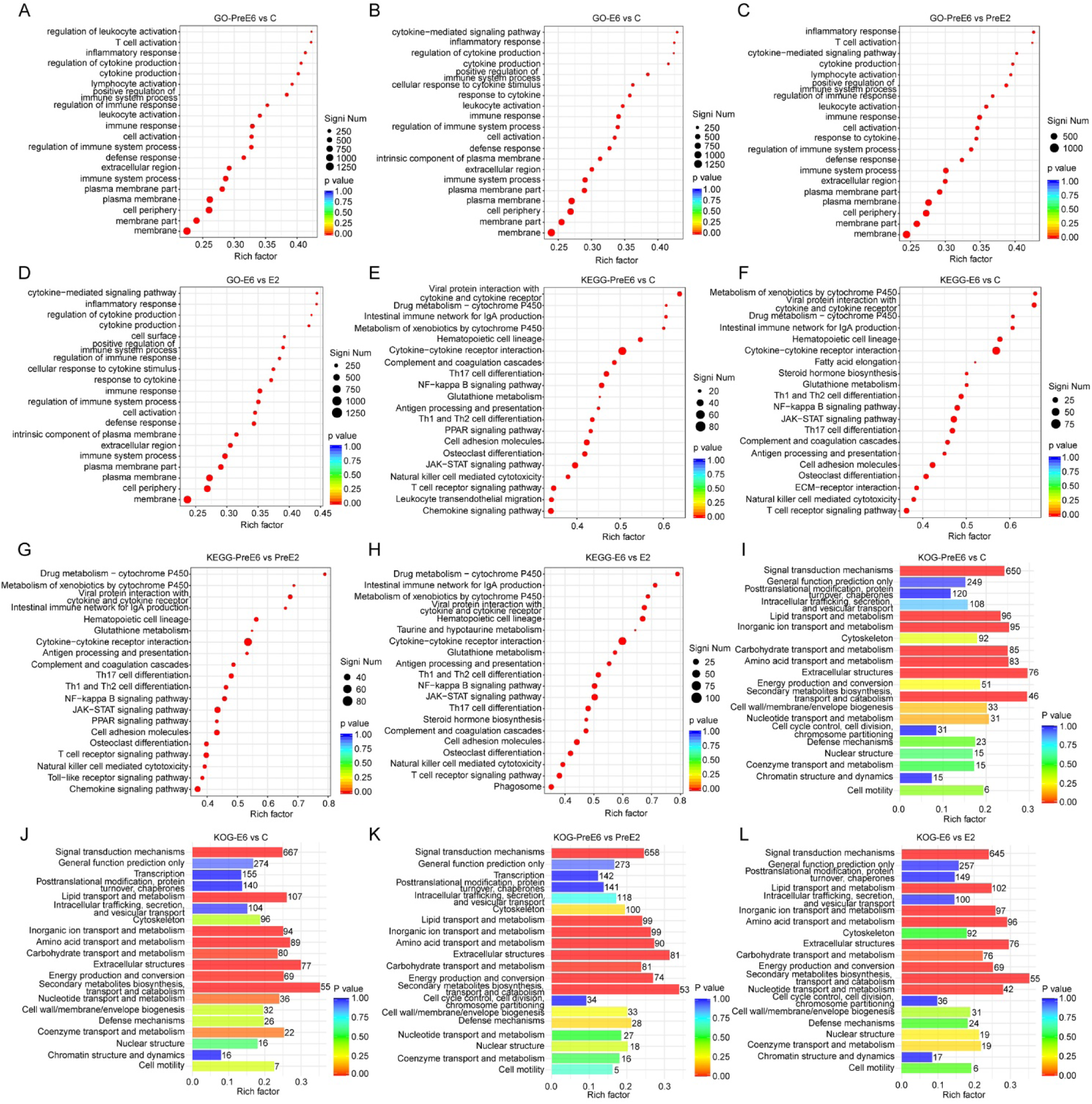
Transcriptome sequencing analysis of chick cecal tissues at different time points after infection *Eimeria* precocious line and parent strain. (A-D) Top 20 enriched GO terms of DEGs in the comparisons of PreE6 vs C, E6 vs C, PreE6 vs PreE2, and E6 vs E2. (E-H) Top 20 enriched KEGG pathways of DEGs in the comparisons of PreE6 vs C, E6 vs C, PreE6 vs PreE2, and E6 vs E2. (I-L) Top 20 enriched KOG classifications of DEGs in the comparisons of PreE6 vs C, E6 vs C, PreE6 vs PreE2, and E6 vs E2. PreE: precocious line oocyst infection; E: parent strain oocyst infection; C: blank control group.

Compared with the transcriptional profiles at 2 dpi, DEGs at 6 dpi in the PreE6 group were uniquely enriched in chemokine signaling, PPAR signaling, and Toll - like receptor signaling pathways, while DEGs in the E6 group were specifically enriched in phagosome, steroid hormone biosynthesis, and taurine and hypotaurine metabolism pathways (Fig. 3G - H). KOG functional classification further supported these transcriptional trends. The DEGs were mainly categorized into extracellular structures, secondary metabolite biosynthesis and metabolism, lipid and amino acid transport and metabolism, and signal transduction mechanisms, indicating functional convergence centered on immune activation and metabolic remodeling (Fig. 3I - L). To fully interpret transcriptional changes at the early infection stage (2 dpi), we displayed the top 20 significantly enriched GO terms, KEGG pathways and KOG classifications of DEGs from PreE2 vs C, E2 vs C, E2 vs PreE2 and E6 vs PreE6 comparisons (Fig. S2)

### Transcriptomic analysis of *Eimeria tenella*

To fully elucidate the linkage between strain developmental differences and host immune responses, differentially expressed genes of parasites within samples were synchronously annotated in this study. Functional enrichment analysis of parasite genes revealed that genes in the P - PreE6 and P - E6 groups were significantly enriched in catabolic processes and cellular component organization, respectively (Fig. 4A - B). Volcano plot analysis identified 271 significantly upregulated genes and 116 significantly downregulated genes in the P - E6 vs P - PreE6 comparison group (Fig. 4C).GO enrichment analysis showed that the parasite-derived DEGs in the P - E6 group, compared with the P - PreE6 group, were significantly downregulated in cellular component terms including membrane, plasma membrane, and extracellular region, and participated in biological processes such as cytokine response, external stimulus response, and type I interferon response (Fig. 4D). KEGG pathway analysis indicated that the DEGs were upregulated in cytokine - cytokine receptor interaction, p53 signaling pathway, RIG - I - like receptor signaling pathway, and biosynthesis-related pathways, while they were downregulated in neuroactive ligand - receptor interaction, absorption - related pathways, and the Hedgehog signaling pathway (Fig. 4E). KOG functional classification further verified that these DEGs were prominently enriched in signal transduction mechanisms and extracellular structure - related categories (Fig. 4F).

**Fig. 4.**
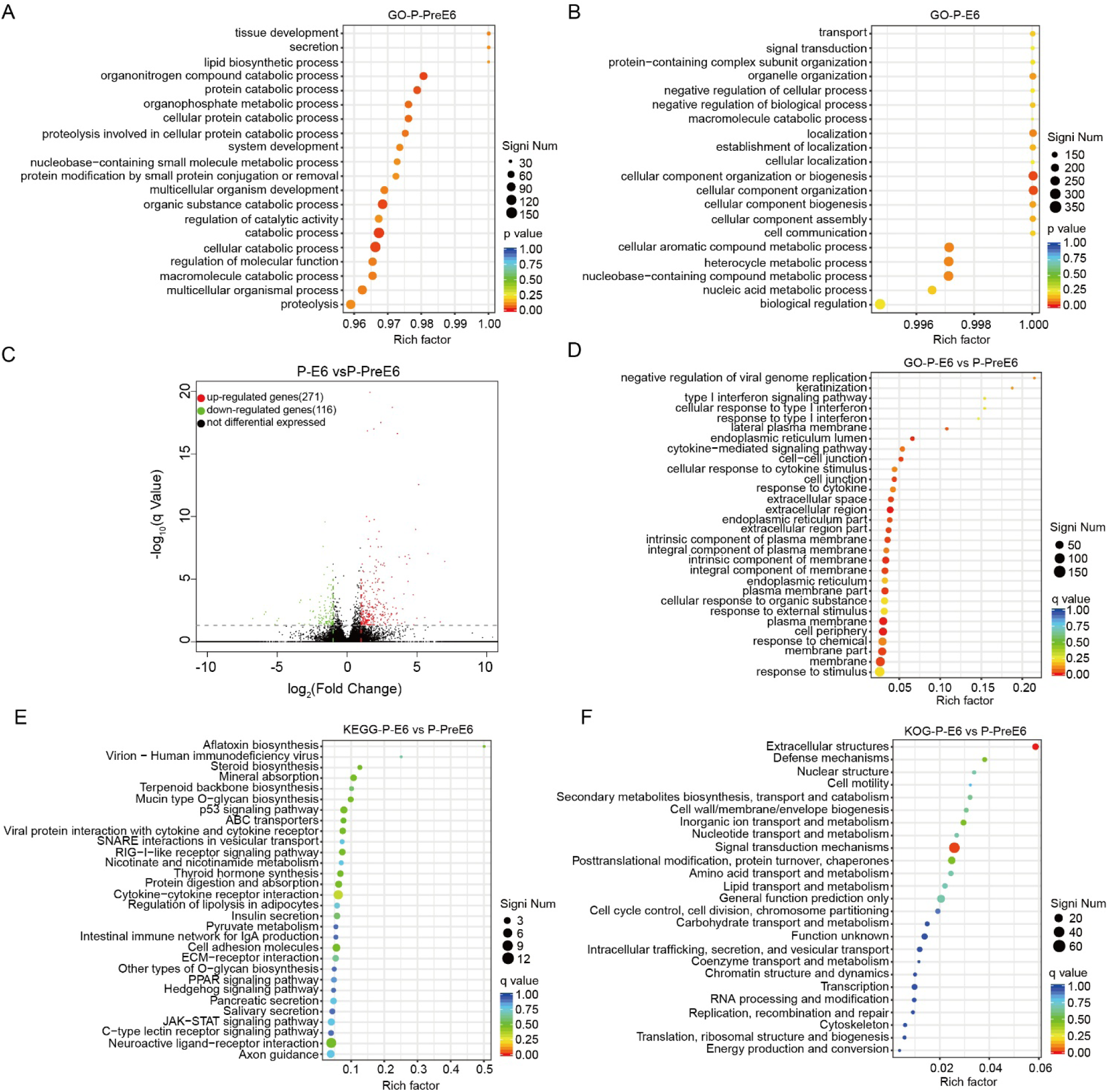
Transcriptomic analysis of *E. tenella* precocious line and parent strain. (A) Top 20 enriched GO terms of parasite genes in the *E. tenella* precocious line. (B) Top 20 enriched GO terms of parasite genes in the *E. tenella* parent strain. (C) Volcano plot showing DEGs between the P-E6 and P-PreE6 groups. (D) Top 30 enriched GO terms of DEGs between the P-E6 and P-PreE6 groups. (E) Top 30 enriched KEGG pathways of DEGs between the P-E6 and P-PreE6 groups. (F) Top 30 enriched KOG classifications of DEGs between the P-E6 and P-PreE6 groups. P-PreE: *E. tenella* precocious line oocysts; P-E: *E. tenella* parent strain oocysts.

### Clinical symptoms and pathological changes of cecal tissues

Gross anatomical observation and histological examination revealed rapid and severe cecal lesions in chicks infected with the parent strain. At 2 dpi, pinpoint hemorrhages and dark brown watery feces were observed in the ceca. Histologically, mild shortening of intestinal villi, shallow crypts, and obvious leukocyte infiltration in the lamina propria were detected. By 6 dpi, bloody cecal cores were formed in the intestinal lumen, accompanied by enlarged and blackened ceca with the intestinal wall thickened 2- 3 times. Severe pan-necrosis and exfoliation of villous epithelium, massive oocyst accumulation in crypts and lamina propria, and extensive hemorrhage were observed in parent strain - infected tissues.

In contrast, the precocious line induced only mild pathological lesions. At 2 dpi, focal mild epithelial necrosis, mucosal hyperplasia, and intensive leukocyte infiltration in the lamina propria were observed. At 6 dpi, only moderate villus shortening and gametocyte - stage oocysts were detected, without diffuse hemorrhage or destruction of intestinal structural integrity (Fig. 5). In summary, the parent strain exhibited rapid pathogenic progression and severe tissue damage, whereas the precocious line displayed remarkably attenuated virulence and controllable pathological lesions.

**Fig. 5.**
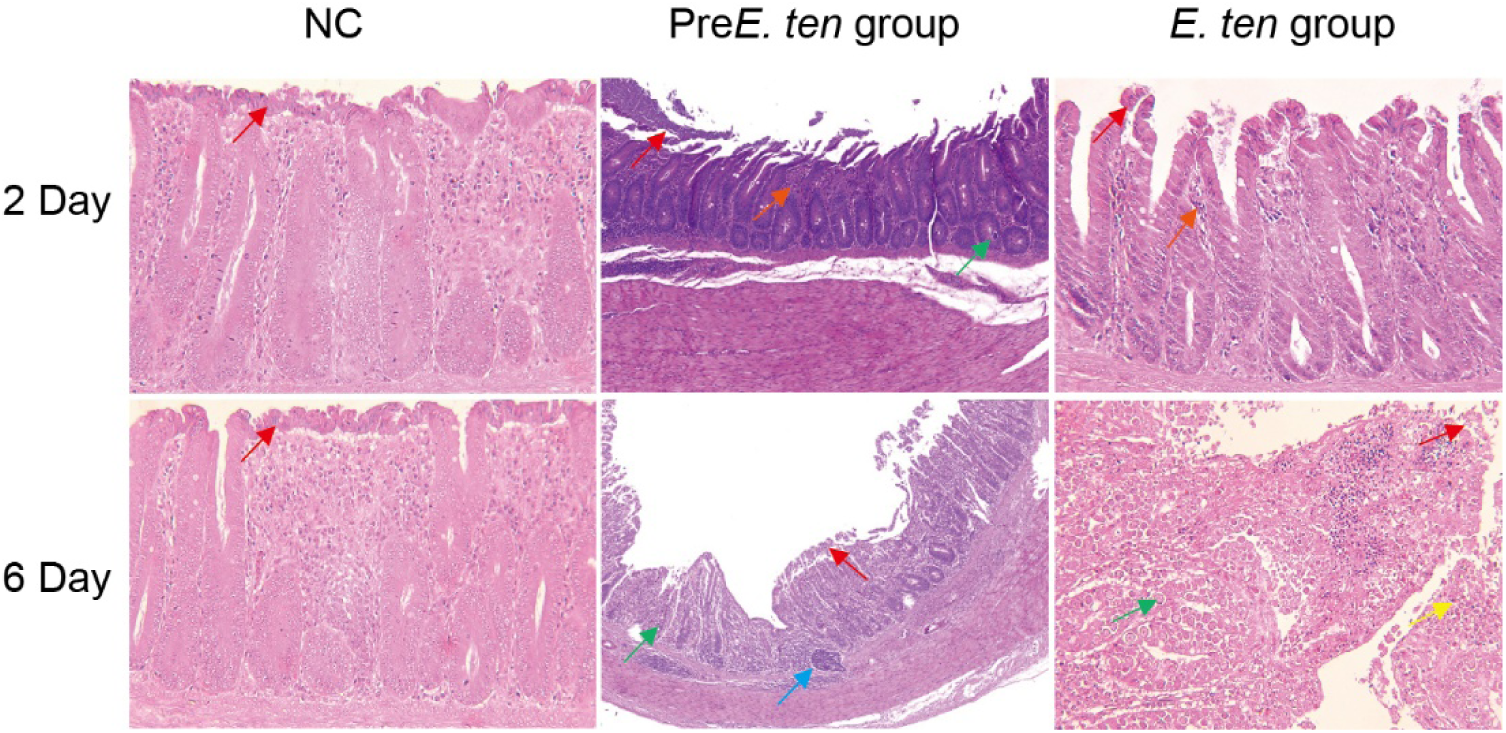
Histopathological sections of chick cecal tissues at 2 and 6 days after infection with different *E tenella* strains. Red arrows: mucosal epithelial cells; orange arrows: inflammatory infiltration; green arrows: coccidial oocysts; blue arrows: lymphoid hyperplasia; yellow arrows: hemorrhage.

### Analysis of local mucosal immune levels in cecal tissues

MTS assay results showed that the proliferative activity of cecal lymphocytes in all infection groups was significantly higher than that in the control group and increased progressively with infection time, with significantly higher proliferation levels at 6 dpi than at 2 dpi. Notably, the lymphocyte proliferation ability in the parent strain-infected group was markedly higher than that in the precocious line group (Fig. S3A). Detection of cecal secretory immunoglobulin A (SIgA) indicated that SIgA levels in both parent and precocious line groups were significantly elevated at 2 dpi compared with the control group, suggesting that mucosal immunity was rapidly activated at the early infection stage. Moreover, SIgA contents were further increased at 6 dpi relative to 2 dpi, demonstrating a continuous enhancement of mucosal immune responses against coccidial infection (Fig. S3B).

### Immune protection assay

No mortality was observed in all immunized groups on day 7 after secondary challenge with 1 × 10^5^ parent strain oocysts (Fig. 6A). Although the body weight gain of all immunized groups remained significantly lower than that of the NC group, it was markedly higher than that of the PC group (Fig. 6B). As shown in Table 4, bloody stool symptoms were still present in all immunized groups, but the bloody stool scores were significantly lower than those in the PC group. Compared with the PC group, chicks pre-immunized with 1 × 10^3^ and 1 × 10^4^ *E. tenella* oocysts (Pre*E. ten* and *E. ten* groups) exhibited significantly reduced cecal OPG values after secondary challenge (Fig. 6C). Immune organ index analysis showed that the thymus indices of the 1 × 10^3^ immunized group and the 1 × 10^4^ precocious line immunized group were higher than that of the PC group. The bursa of Fabricius index was also significantly increased in the 1 × 10^4^ parent strain immunized group, while no significant difference in spleen index was detected among all groups (Fig. 6D). In conclusion, pre - immunization with low - dose precocious line and parent strain oocysts can induce effective immune protection, significantly alleviate intestinal lesions, reduce oocyst shedding and mortality after secondary infection, and exert positive effects on the development of immune organs in chicks.

**Fig. 6.**
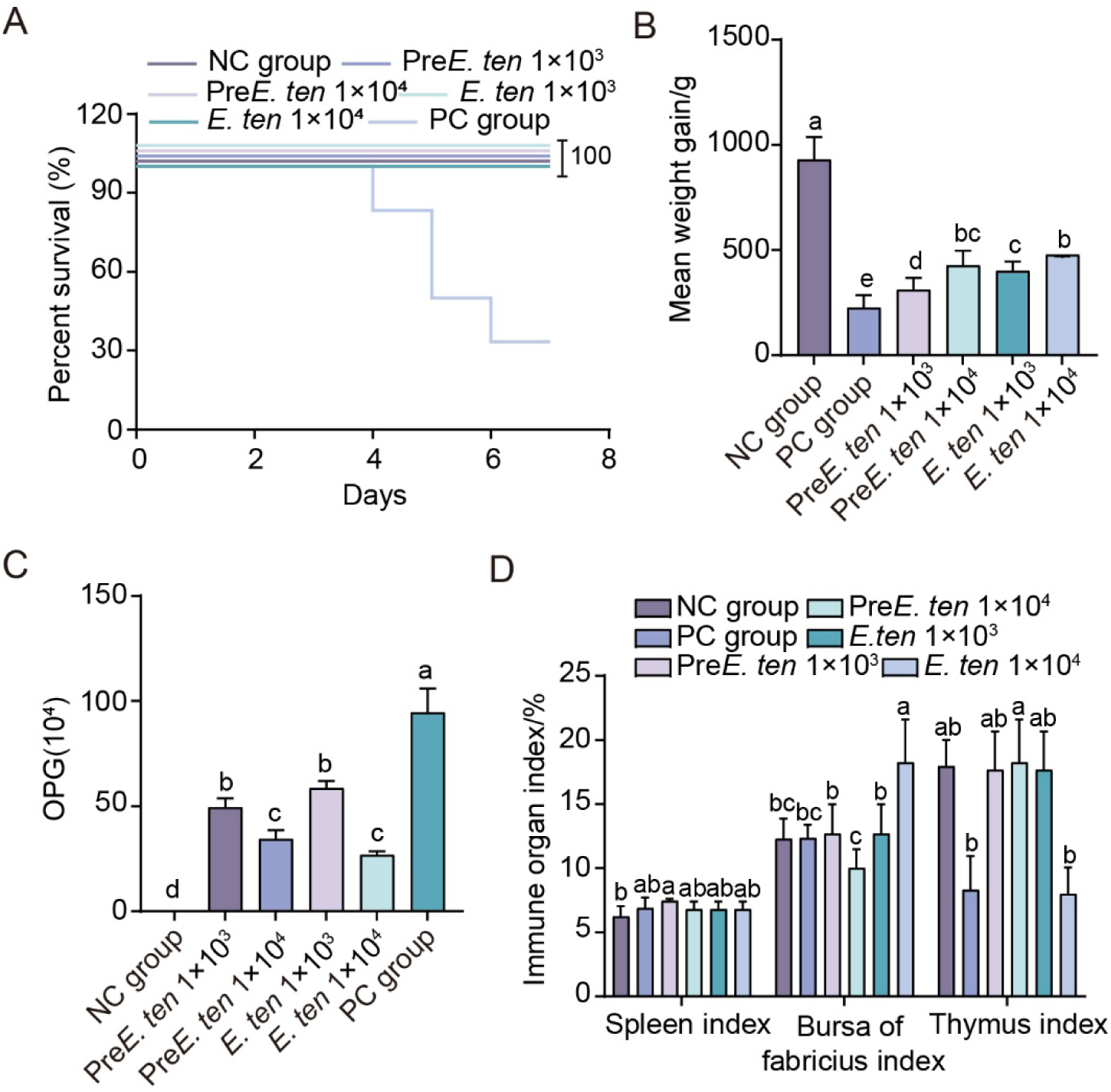
Immune protection assay of sporulated oocysts of *E. tenella* precocious line and parent strain. (A) Effects of different dosages of precocious line and parent strain oocysts on chick survival rate during immune protection detection. (B) Effects of different dosages of precocious line and parent strain oocysts on chick body weight gain during immune protection detection. (C) OPG of cecal contents in each group in the immune protection assay. (D) Changes of immune organ indices in each group. Values marked with different lowercase superscript letters (a–d) within the same column indicate significant differences at *P* < 0.05.

**Table 4.** Statistical analysis of bloody stool scores after secondary infection.

| Group | Oocyst infection dose | 84 h | 93 h | 108 h | 117 h | 132 h | 141 h | 156 h | 165 h | Score of bloody feces |
| --- | --- | --- | --- | --- | --- | --- | --- | --- | --- | --- |
| NC | 0 | 0 | 0 | 0 | 0 | 0 | 0 | 0 | 0 | 0<sup>c</sup> |
| PC | - | 4 | 4 | 4 | 3 | 3 | 2 | 0 | 0 | 2.5±1.69<sup>a</sup> |
| Pre*E. ten* | 1.0×10<sup>3</sup> | 2 | 3 | 2 | 1 | 1 | 0 | 0 | 0 | 1.13±1.13<sup>b</sup> |
| Pre*E. ten* | 1.0×10<sup>4</sup> | 0 | 2 | 1 | 0 | 1 | 0 | 0 | 0 | 0.5±0.76<sup>bc</sup> |
| *E. ten* | 1.0×10<sup>3</sup> | 1 | 3 | 1 | 1 | 0 | 0 | 0 | 0 | 0.75±1.04<sup>bc</sup> |
| *E. ten* | 1.0×10<sup>4</sup> | 0 | 1 | 0 | 0 | 0 | 0 | 0 | 0 | 0.13±0.35<sup>bc</sup> |

Values marked with different lowercase superscript letters (a-d) within the same column indicate significant differences at $P < 0.05$ .

## Discussion

In the present study, temporal transcriptomic profiling of chick cecal tissues was performed at two critical developmental stages of *E. tenella*, schizogony (2 dpi) and gametogony (6 dpi), to systematically compare the differential host molecular responses induced by virulent parent strain and attenuated precocious line infections. Transcriptomic results revealed that only approximately 200 DEGs were identified at 2 dpi, indicating a mild transcriptional remodeling of cecal tissues at the early infection stage. In contrast, the number of DEGs increased dramatically to 2000∼3500 at the gametogony stage (6 dpi), demonstrating that gametogony serves as the core activation stage for cecal mucosal immune responses in chicks. Accordingly, subsequent functional enrichment analyses in this study were primarily focused on transcriptomic data obtained at 6 dpi. Although both strains initiated the upstream cytokine - JAK/STAT innate immune pathway, their downstream signaling branches exhibited distinct divergence, mediating two completely opposite regulatory programs corresponding to destructive inflammation and balanced immune homeostasis, respectively. Both the parent strain and precocious line activated the cytokine - JAK/STAT innate immune axis, whereas their downstream regulatory trajectories were entirely differentiated, forming two distinct molecular patterns: uncontrolled inflammation leading to tissue damage and balanced inflammation facilitating mucosal repair. The parent strain triggered persistent excessive activation of the NF - κB signaling pathway, which synchronously drove three destructive cascades, including coagulation - complement - ECM positive feedback loops, steroid metabolic disturbance, and antioxidant system exhaustion, ultimately resulting in severe organic lesions in the cecum. Previous studies have confirmed that blocking the NF - κB pathway effectively alleviates *Eimeria* - induced intestinal injury, further verifying the pivotal pathogenic role of NF - κB signaling in coccidiosis (Meng et al., 2024). Sustained hyperactivation of NF - κB promotes the massive release of pro - inflammatory mediators and upregulates tissue factors to initiate the extrinsic coagulation pathway. Complement activation generates C3a and C5a, which recruit inflammatory infiltrating cells, while coagulation products in turn amplify complement activation, thereby forming a bidirectional inflammation - thrombosis amplification loop. This cascade continuously degrades the extracellular matrix and disrupts intestinal epithelial barrier integrity, which is consistent with previous findings regarding EtROP17 - mediated cellular injury and inflammatory bowel disease-related mechanisms (Scaldaferri et al., 2011; Meng et al., 2022; Pryzdial et al., 2022).

Furthermore, severe infection induces systemic stress disorders and abnormal activation of the hypothalamic - pituitary - adrenal axis. Excessive steroid secretion suppresses Th1/Th17 protective immune responses and impairs host parasite clearance capacity. Meanwhile, persistent oxidative stress induces massive taurine consumption, leading to antioxidant system exhaustion and reactive oxygen species accumulation, which further accelerates intestinal epithelial necrosis (Raber, 1998; XU et al., 2015; Seneff et al., 2025). The pathological manifestations of pinpoint cecal hemorrhage at 2 dpi and extensive cecal core formation with mucosal exfoliation at 6 dpi are the intuitive morphological outcomes of this progressive pathological cascade.

In sharp contrast to the destructive regulatory pattern of the parent strain, the precocious line specifically activates PPAR signaling, TLR signaling, and leukocyte transendothelial migration pathways, constructing a complete regulatory cascade of “immune sensing - inflammatory restriction - pathogen elimination - tissue repair” that precisely balances anti - parasite immune defense and intestinal homeostasis. PPAR - γ represents the most distinctive core molecular feature distinguishing the precocious line from the parent strain, which is significantly enriched as early as 2 dpi. By binding to the NF - κB p65 subunit and upregulating IκBα expression, PPAR - γ blocks NF - κB nuclear translocation and inhibits the excessive secretion of pro-inflammatory factors such as TNF - α and IL - 6, exerting an early lipid - mediated anti - inflammatory effect (Ju et al., 2020). The TLR signaling pathway is activated in a precise temporal pattern during precocious line infection: chTLR7/15 is activated at the early stage for immune initiation, while chTLR21 sustains continuous immune signaling. This temporal activation synchronously induces chemokine-dependent directional leukocyte migration to infection sites for parasite elimination, accompanied by the secretion of anti-inflammatory cytokines (IL - 10, TGF - β) to facilitate mucosal repair and avoid cytokine storm caused by excessive TLR activation (Wang et al., 2024). This refined regulatory mechanism only induces mild local mucosal damage without diffuse hemorrhage or full - thickness epithelial necrosis.

Based on the comprehensive comparison of differential pathway activation, an integrated mechanistic model illustrating the divergent pathogenic mechanisms of the parent strain and precocious line was established in this study (Fig. 7). Both strains share the upstream NF - κB - dependent innate immune initiation signal. However, the precocious line possesses an early PPAR - γ-mediated anti-inflammatory regulatory module that negatively constrains excessive NF - κB activation. In contrast, the parent line lacks this negative regulatory mechanism, and persistent NF - κB hyperactivity drives multiple pathological cascades, ultimately leading to complete collapse of intestinal structural integrity. These two opposite regulatory modes constitute a distinct bipolar immune regulatory paradigm during *E. tenella* infection.

**Fig. 7.**
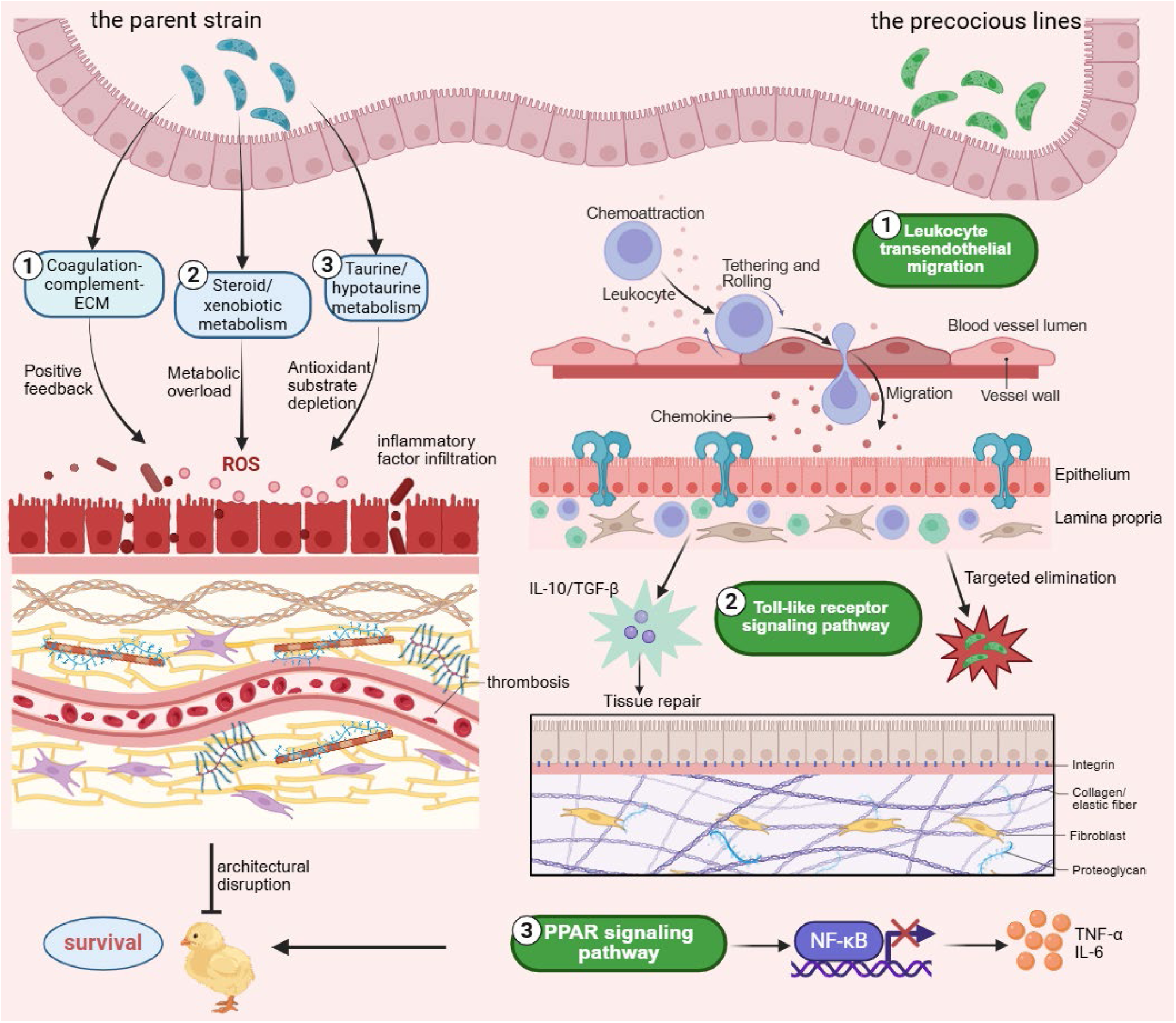
Schematic diagram of the differential pathogenic mechanisms in chicks infected with *E. tenella* precocious line and parent strain oocysts. The parent strain synchronously triggers three severe damaging effects including coagulation and thrombosis formation, metabolic disorders, and antioxidant system exhaustion, ultimately leading to complete collapse of intestinal structural integrity. In contrast, the precocious line early activates the PPAR-γ signaling pathway to negatively inhibit excessive NF-κB activation, accompanied by accurate immune recognition via the TLR pathway and targeted leukocyte recruitment and infiltration. This regulatory pattern efficiently eliminates parasites while restricting inflammatory responses within a safe and controllable range, forming a distinct bipolar regulatory contrast between the two *E. tenella* strain.

Although precocious attenuated strains have been commercially applied for avian coccidiosis prevention and control, most current studies focus on parasite-derived virulence genes, lacking systematic temporal profiling of host mucosal immune responses across schizogony and gametogony stages. Based on dynamic transcriptomic data at 2 dpi and 6 dpi, the present study identified PPAR-γ as the core hub for balancing intestinal inflammatory homeostasis and fully clarified the molecular mechanisms underlying uncontrolled inflammation induced by the parent strain and controllable inflammation triggered by the precocious line, filling the research gap in host - parasite temporal interaction mechanisms during *E. tenella* infection.

The NF - κB - mediated tissue injury pathway and PPAR - γ - dominated anti -inflammatory and mucosal repair network elucidated in this study systematically reveal the intrinsic molecular basis of the low virulence and high immunogenicity of *E. tenella* precocious lines. These findings provide comprehensive theoretical support for targeted intervention against poultry intestinal inflammation, optimization of evaluation systems for attenuated strain screening, and exploration of novel targets for antibiotic - free green prevention and control of avian coccidiosis.

## Conclusion

Based on comparative temporal transcriptomic analysis of chick cecal tissues at 2 dpi and 6 dpi, the present study confirmed that the pathogenic discrepancy between *E. tenella* parent strain and precocious line is determined by the bipolar divergence of host inflammatory regulatory pathways. The virulent parent strain induces persistent NF - κB hyperactivation, triggering a chained pathological cascade that causes irreversible cecal tissue damage. In contrast, the precocious line significantly upregulates PPAR - γ signaling at the early infection stage, which cooperates with the TLR pathway to precisely balance anti-parasite immune defense and intestinal homeostasis. PPAR - γ is identified as the core molecular hub mediating mild tissue damage and robust mucosal immune responses during precocious line infection. Currently, this precocious attenuated strain has been commercially applied for the development of live - attenuated coccidiosis vaccines. This study systematically elucidates the unique molecular regulatory mechanism of mucosal immunity triggered by the precocious line, improves the theoretical system for the pathogenic characteristics of attenuated *E. tenella* strains, and provides critical molecular evidence for targeted antibiotic-free prevention and control of avian coccidiosis.

## Author contributions

WM: Writing-Original Draft, Investigation, Formal Analysis. KD: Investigation, Data curation. TY: Investigation, Validation. XL: Investigation, Validation. SN: Investigation. MD:Validation. XL: Supervision. JA: Investigation. DY: Supervision, Conceptualization, Writing - review & editing. QL: Project administration, Supervision, Writing - review & editing.

## Funding

This work was supported by the Key Research and Development & Transformation Special Project of Xizang Autonomous Region Science and Technology Program (Grant No. XZ202601ZY0136-3); the Base and Talent Program of Xizang Autonomous Region Science and Technology Program (Grant No. XZ202502JD0028); the CSC Scholarship Program for Global Rural Revitalization Talents in Animal Health and Husbandry (XCZXRC20230006); and the Young Teacher Research Innovation Promotion Program of Beijing University of Agriculture (Grant No. QJKC-2023003).

## Conflict of interest

The author(s) declared that this work was conducted in the absence of any commercial or financial relationships that could be construed as a potential conflict of interest.

## Generative AI statement

Generative AI (Doubao) was used solely for language polishing (grammar, clarity, and style only) during manuscript preparation. The authors have thoroughly reviewed and verified all content, take full responsibility for the manuscript, and confirm that AI was not used to generate data, perform analyses, or draw scientific conclusions.

## Publisher’s note

All claims expressed in this article are solely those of the authors and do not necessarily represent those of their affiliated organiza tions, or those of the publisher, the editors and the reviewers. Any product that may be evaluated in this article, or claim that may be made by its manufacturer, is not guaranteed or endorsed by the publisher.

## Figure captions

**Supplementary Figure 1.**
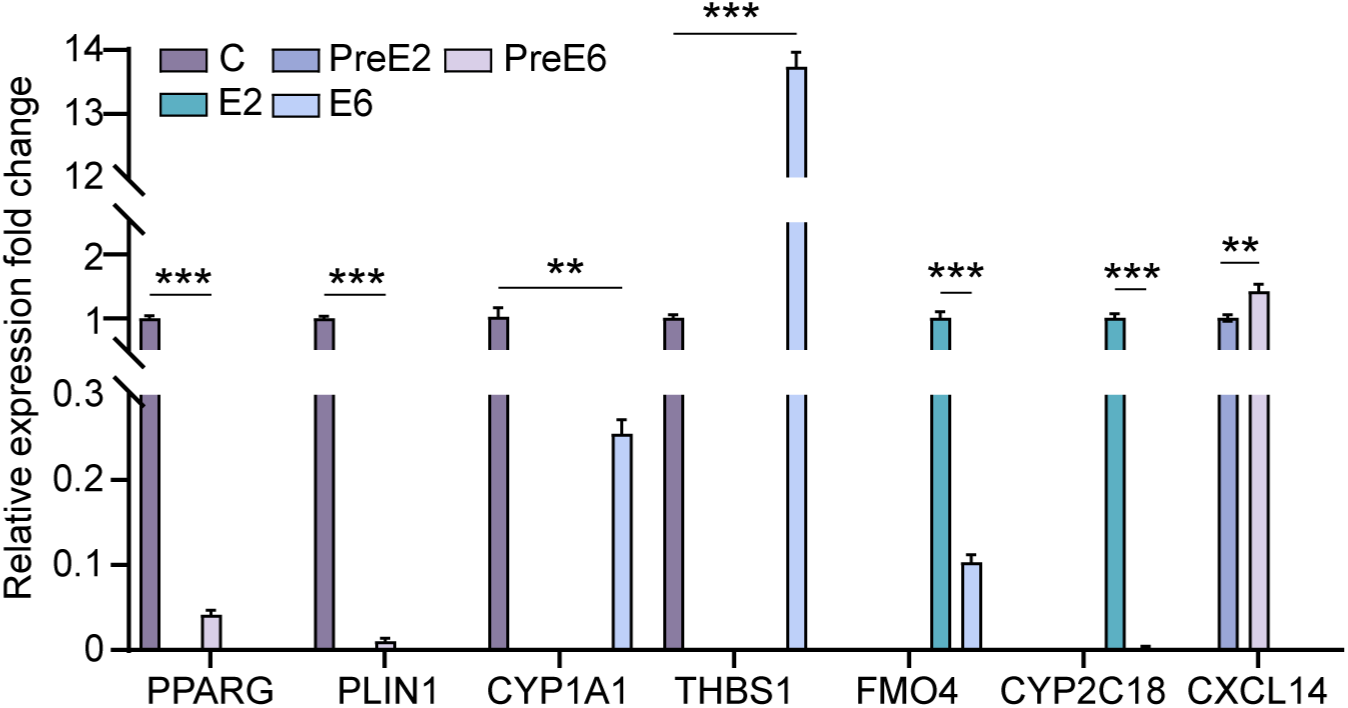
QRT-PCR validation of seven key differentially expressed genes. The validated genes were screened from different comparison groups: PPARG and PLIN1 in PreE6 vs C; CYP1A1 and THBS1 in E6 vs C; FMO4 and CYP2C18 in E6 vs E2; and CXCL14 in PreE6 vs PreE2. C: blank control group; PreE2: precocious line infection group at 2 dpi; PreE6: precocious line infection group at 6 dpi; E2: parent strain infection group at 2 dpi; E6: parent strain infection group at 6 dpi. \*\**P* < 0.01, \*\*\**P* < 0.001.

**Supplementary Figure 2.**
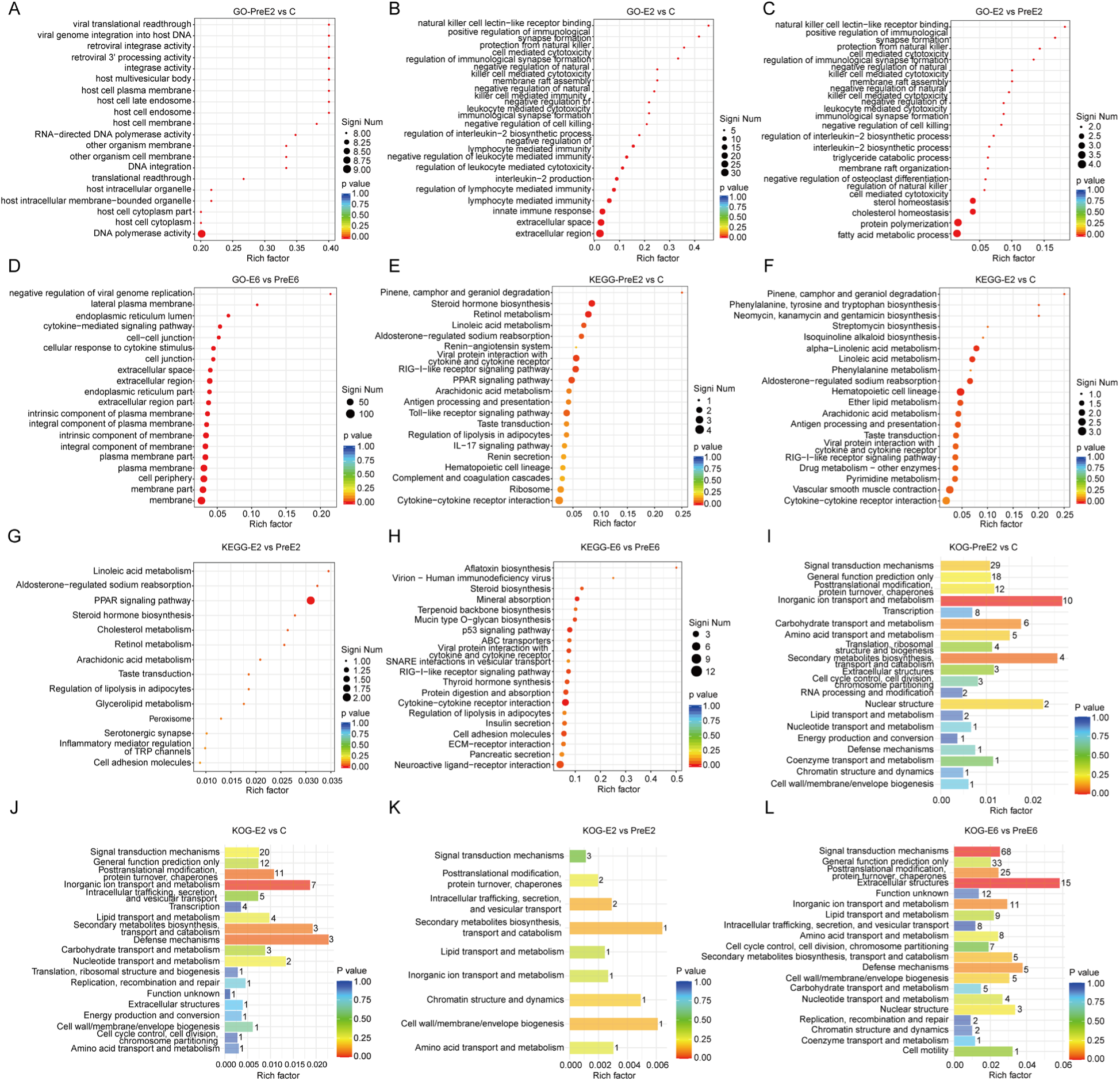
Transcriptome sequencing analysis of chicken cecal tissues at different time points after infection with precocious and parent strains of *E. tenella*. (A-D) Top 20 enriched GO terms of DEGs in comparisons of PreE2 vs C, E2 vs C, E2 vs PreE2, and E6 vs PreE6. (E-H) Top 20 enriched KEGG pathways of DEGs in comparisons of PreE2 vs C, E2 vs C, E2 vs PreE2, and E6 vs PreE6. (I-L) Top 20 KOG classifications of DEGs in comparisons of PreE2 vs C, E2 vs C, E2 vs PreE2, and E6 vs PreE6. PreE, precocious line oocysts; E, parent strain oocysts; C, uninfected control group.

**Supplementary Figure 3.**
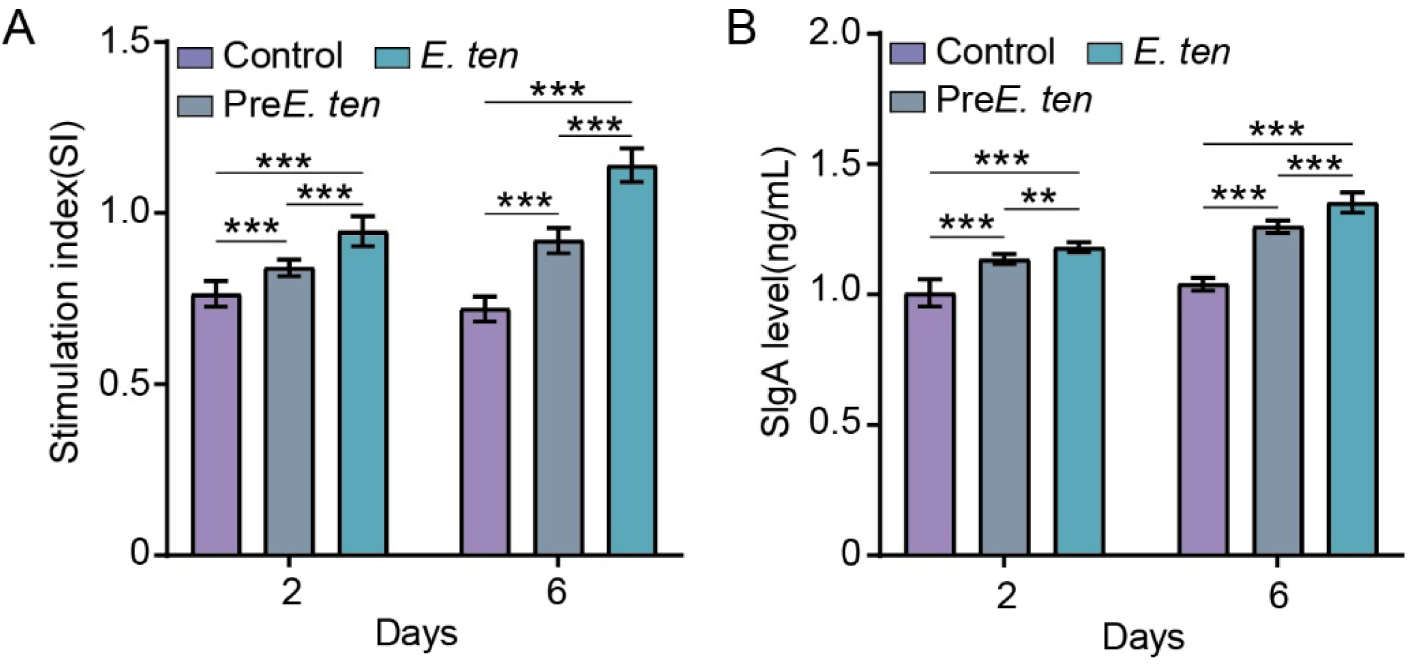
Detection of immune indicators in chick cecal tissues at 2 and 6 days post infection with different virulent *E. tenella* oocysts. (A) Stimulation index (SI) of lymphocyte proliferation; (B) De

## Notes

### Competing Interest Statement

The authors have declared no competing interest.

